# Integrating transcriptomic data with metabolic model unravels the electron transfer mechanisms of *Methanosarcina barkeri*

**DOI:** 10.1101/2025.01.27.635023

**Authors:** Wentao Tang, Sen Lin, Yangfan Deng, Gang Guo, Guanghao Chen, Tianwei Hao

## Abstract

Methanogenic archaea, particularly *Methanosarcina*, are pivotal to the global carbon cycle and renewable energy production due to their versatile metabolic capabilities. Although transcriptomic analysis is widely employed to identify key genes and pathways in *Methanosarcina* under various methanogenic conditions—including the emerging direct interspecies electron transfer (DIET)-based methanogenesis—the weak correlation between gene expression levels and protein abundances poses challenges for interpreting transcriptomic data. To address this, we integrated transcriptomic data into a metabolic model of *Methanosarcina barkeri* for the first time, enabling more refined predictions and enhancing the interpretability of transcriptomic insights. This novel integrated model was subsequently utilized to simulate aceticlastic, hydrogenotrophic, and DIET-based methanogenesis. The results revealed that previous assumptions failed to account for the role of the CO₂ reduction pathway in aceticlastic methanogenesis. The model also successfully captured key transcriptomic features of DIET-based methanogenesis, clarifying the functional roles of crucial enzymes like the F_420_H_2_ dehydrogenase Fpo and the transmembrane hydrogenase Vht in electron transfer. This integrative approach provided a deeper understanding of electron transfer mechanisms in *M. barkeri* and offered valuable insights for advancing methanogen-based biotechnologies. Moreover, the study critically evaluated the relationship between gene expression and metabolic flux, establishing a practical framework for deriving meaningful insights from the growing volume of transcriptomic data.

## 1. Introduction

Methanogenic archaea produce approximately one billion tons of methane annually on a global scale [1]. About two-thirds of this biogenic methane is originated from acetate cleavage via a process called aceticlastic methanogenesis, while the remaining one-third is generated through hydrogenotrophic methanogenesis, where carbon dioxide (CO_2_) is reduced to methane using electrons derived from hydrogen (H_2_) [1, 2]. Currently, only two methanogenic genera, *Methanosarcina* and *Methanothrix* (formerly *Methanosaeta*), are identified as capable of using acetate for methane production [3]. Unlike *Methanothrix*, which exclusively metabolizes acetate, *Methanosarcina* can utilize a broader range of substrates, including acetate, H_2_/CO_2_, and methyl compounds [4, 5], thereby enhancing its adaptability in diverse environments [6, 7]. Furthermore, *Methanosarcina* species have recently been demonstrated to actively participate in direct interspecies electron transfer (DIET) with *Geobacter metallireducens*, a well-known model microorganism that serves as an electron donor during DIET [8, 9]. DIET represents a form of syntrophic metabolism where microorganisms exchange electrons through direct electrical connections, in contrast to mediated interspecies electron transfer (MIET), which relies on diffusible electron mediators such as H_2_ or, occasionally, formate [10, 11]. The discovery of DIET has revolutionized our understanding of methanogenesis, as it allows *Methanosarcina* to simultaneously generate methane from acetate cleavage and CO_2_ reduction by extracellular electrons, thus enhancing methane yield [8, 12]. This finding underscores the ecological and biotechnological significance of *Methanosarcina* in methane generation and highlights the need to elucidate their cellular metabolism under different methanogenic conditions [13, 14]. To date, the research on *Methanosarcina*’s cellular metabolism has primarily focused on understanding how electrons are utilized by electron acceptors (e.g., CO_2_, methyl group of acetyl-CoA) to produce methane [1, 15, 16]. However, the mechanisms underlying electron transfer from electron donors (e.g., H_2_, carbonyl group of acetyl-CoA, DIET electron donor) to electron acceptors, as well as the processes of energy generation during electron transfer, particularly in DIET-based methanogenesis, remain unresolved [9, 17, 18].

Transcriptomic analysis is a powerful tool for understanding cellular responses to various conditions, including aceticlastic, MIET-based, and DIET-based methanogenesis [9, 12, 19, 20]. Typically, statistical methods are applied to identify differentially expressed genes that may signal functional changes. However, metabolic activities are more directly tied to protein levels, and gene expression levels often exhibit weak correlation with protein abundances, with correlation coefficients typically below 0.5 [21]. This weak correlation is attributed to factors like post-transcriptional modifications, differing transcript and protein half-lives, and measurement noise [22]. As a result, transcriptomics alone is insufficient to fully capture metabolic activities under varying methanogenic conditions [19]. Moreover, transcriptomics also lacks metabolic network context, failing to consider pathway topology, regulation, and interactions that govern metabolic activities [23]. For instance, a comparative transcriptomic study on DIET and MIET proposed that the upregulated enzyme F_420_: phenazine oxidoreductase (Fpo) functions to reduce coenzyme F_420_ while oxidizing reduced methanophenazine (MPH_2_) during DIET [18]. It was hypothesized that the reduced coenzyme F_420_H_2_ then supplies electrons for producing reduced ferredoxin (Fd_red_) via the electron-bifurcating heterodisulfide reductase complex (HdrABC), thus facilitating methane generation from CO_2_ reduction. However, this assumption conflicts with the role of aceticlastic methanogenesis, which co-occurs with CO_2_ reduction during DIET and requires Fpo to function in reverse—oxidizing F_420_H_2_ and reducing methanophenazine (MP) [8, 24].

Furthermore, a subsequent transcriptomic study by the same group revealed that several genes encoding Fpo and HdrABC were not expressed during DIET [9], contradicting their earlier study [18]. Transcriptomic data alone cannot resolve such contradictions, as it lacks information on reaction directionality and cannot distinguish enzymes with opposing effects on reaction flux, highlighting the necessity for additional information to accurately interpret expression changes [25].

Integrating transcriptomic data with genome-scale metabolic models (GEMs) offers a robust solution by placing gene expression within the context of metabolic networks, thereby enhancing our understanding of the functional implications of expression changes [26, 27]. GEMs are computational reconstructions of strain-specific metabolic networks that incorporate information on metabolites, reactions, pathways, and their associated genes of specific organisms [28]. These models provide a predefined biochemical reaction network constrained by mass balance and thermodynamics, which can compensate for the noise and variability inherent in transcriptomic data, thus enabling objective interpretation of expression data [28, 29]. Meanwhile, expression data offers condition-specific context that can narrow down the solution space, thereby enhancing the accuracy and specificity of simulations [26, 30]. This integrated approach has only been applied to a few well-investigated model microbes, including *Escherichia coli*, *Mycobacterium tuberculosis*, and *Saccharomyces cerevisiae* [26, 29, 31]. Despite advancements in next-generation sequencing technologies generating entensive genomic and transcriptomic data, the construction of curated, strain-specific GEMs has lagged behind. This is due to the fact that the construction of high-quality GEMs is a complex and time-intensive process involving 96 steps, as outlined in the standard protocol by Thiele and Palsson [32]. To achieve more efficient simulations, the core metabolic model approach was developed in our previous study [33], which simplifies GEM construction by focusing on a subset of biologically significant and well-characterized pathways familiar to readers with foundational biochemistry knowledge [33, 34]. This simplification reduces computational demands while maintaining the accuracy and utility of metabolic models. The core metabolic model concept is particularly suitable for anaerobic microbes, such as methanogens, where the majority of the carbon and energy sources are allocated to energy metabolism (e.g., methanogenesis) [35, 36]. Consequently, the biosynthetic pathways become less critical, allowing them to be simplified for modelling purposes.

Therefore, to investigate the electron transfer mechanisms underlying different methanogenic conditions, the present study developed a core metabolic model for the model organism *Methanosarcina barkeri* MS and integrated it with the available transcriptomic data to improve model predictions. Simulations were conducted for three modes of methanogenesis—DIET, MIET, and aceticlastic—to obtain comprehensive insights into the electron transfer mechanisms of *M. barkeri*, which could improve our understanding of the methanogenesis process and contribute to optimizing its performance under various conditions.

## 2. Materials and methods

### 2.1. Model development

In this study, a draft metabolic model for *M. barkeri* was first generated using the ModelSEED pipeline [37], based on the annotated genome of *M. barkeri* MS (NCBI accession number: GCA_000970025.1). The genome was uploaded to the ModelSEED server, where genes were automatically mapped to reactions using its internal database. The resulting draft model included gene-protein-reaction (GPR) associations, a list of reactions, and associated metabolites. Subsequently, a core metabolic model of *M. barkeri* was derived by selecting essential reactions representing central carbon and energy metabolism [33]. Specifically, core reactions related to glycolysis, tricarboxylic acid (TCA) cycle, methanogenesis, and electron transport chain were retained (Fig.S1), while most other reactions were excluded unless they were essential for biomass synthesis. The desired output of the metabolic model was defined by an objective function, typically aimed at maximizing biomass production or the synthesis of a biotechnologically important metabolite. During flux balance analysis (FBA), this objective function is optimized to predict the contribution of each reaction to the overall metabolic goal. In this study, the biomass objective function was established based on biomass precursors (stoichiometry obtained from [34]) rather than macromolecules such as proteins, RNA, lipid, DNA (Fig.S1), with growth-associated ATP maintenance was set at 65.18 mmol/gDW [38].

The core model was manually refined and curated to enhance its accuracy and reliability, following the protocol described by Thiele and Palsson [32]. Reactions, metabolites, and GPR associations were systematically checked against existing literature, expert knowledge, and experimental data specific to *M. barkeri*. Unique reactions, particularly those involving proton transport, were carefully curated to reflect the organism’s metabolism. For example, the Na^+^/H^+^ antiporter (Nha) was configured to exchange one sodium ion per proton [39]. Standard Gibbs free energy information was incorporated, and an electron uptake reaction for DIET was included. A detailed spreadsheet containing all reaction formulas, metabolite information, thermodynamic data, and GPR associations is available in the supplementary Excel file.

### 2.2. Model simulation using flux balance analysis (FBA)

The metabolic model was transformed into a stoichiometric matrix using the MATLAB Cobra toolbox [40], where rows represent metabolites and columns represent reactions. Simulations were performed using FBA, a constraint-based approach for calculating the flow of metabolites through the metabolic network [41]. The metabolic model was simulated under the assumption of steady-state condition, where the flux through all internal reactions balances the rates of substrate uptake and product formation. FBA determines the optimal distribution of metabolic flux that maximizes a defined objective function, typically biomass production, while adhering to the constraints prescribed by the stoichiometric matrix, mass balance, and growth medium. This approach provides in silico growth predictions and insights into optimal metabolic pathways.

Three modes of methanogenesis—DIET, MIET, and aceticlastic—were simulated in this study with the objective function of maximizing biomass synthesis. Constrains were applied based on limited uptake rates of extracellular electron (DIET), hydrogen (MIET), and acetate (aceticlastic methanogenesis). Additionally, to obtain insights into the metabolic flexibility and robustness, flux variability analysis (FVA) was performed to explore the feasible ranges of reaction fluxes that meet the original objective, while considering an optimality factor [42].

### 2.3. Integration with transcriptomic data

Transcriptomic data was then utilized to refine the metabolic model by mapping gene expression data directly onto metabolic reactions. Reactions were considered inactive if the associated genes were not expressed, and potentially active if gene expression was detected [43]. RNA sequencing datasets used for this purpose were retrieved from the NCBI Sequence Read Archive (SRA) and processed using the DOE’s Systems Biology Knowledgebase (KBase) platform (http://kbase.us). The accession numbers of sequencing data under different methanogenic conditions were as follow: aceticlastic (SRX5544684), DIET (SRX4966420, SRX4966418, and SRX4966417), MIET (SRX4966422, SRX4966421, and SRX4966419), and DIET-2 (SRX14923653). Sequencing data quality was assessed using FASTQC [44]. Subsequently, reads were aligned to the genome of *M. barkeri* MS (GCA_000970025.1) using HISAT2 [45]. The aligned reads were then assembled with StringTie [46] to quantify normalized gene expression levels in TPM (Reads Per Kilobase Million).

Expression value was assigned to gene-associated reactions based on Boolean gene-protein-reaction (GPR) rules. It is crucial to acknowledge that not all genes had a one-to-one mapping with reactions, as multiple genes were often associated with a single reaction. For reactions catalysed by enzyme complexes, the minimum expression level among all the subunits was chosen. For instance, reaction HDR is catalysed by heterodisulfide reductase (HdrDE) which consists of two subunits, D and E, with GPR association of *MSBRM_RS15135 and MSBRM_RS15140*. During aceticlastic methanogenesis, these subunits showed TPM values of 6.34 and 6.10, respectively, and the minimum value (6.10) was selected to represent the expression level of the HDR reaction. In contrast, for reactions catalysed by isoenzymes or multiple copies of an enzyme, the expression levels of these associated genes were summed. For example, reaction 04042 is catalysed by two copies of ADP-dependent glucokinase and associated with GPR rule of *MSBRM_RS13255 or MSBRM_RS13250*. In this scenario, the transcriptional levels of both genes were summed to represent the expression level of the reaction. The gene expression data and model-predicted flux from FBA analysis were visualized using Escher map [47], providing a contextualized view of multiple datasets within the metabolic network context under various methanogenic conditions.

## 3. Results and discussion

### 3.1. Metabolic modelling of aceticlastic methanogenesis

A core metabolic model of *M. barkeri* was first developed, incorporating the activity of 204 metabolic genes, 117 metabolites, and 101 reactions based on the latest literature evidence, expert insights, and experimental data. To validate the predictive accuracy of the model, simulation of aceticlastic methanogenesis via FBA analysis was performed and compared with published experimental data. As expected, the model successfully predicted that *M. barkeri* could sustain growth using acetate as its sole carbon and energy source. The predicted growth rate and yield aligned with experimental data collected from multiple literature sources (Fig. 1) [36, 48–50]. The predicted metabolic fluxes of central carbon and energy metabolic pathways of aceticlastic methanogenesis were illustrated in Fig. 2A. After activation of acetate into acetyl-CoA (via ACK and PTA reactions in Fig. 2A), a significant majority (over 97%) of acetyl-CoA was predicted to be utilized for energy metabolism (i.e., methanogenesis) through the carbon monoxide dehydrogenase/acetyl-CoA synthase (CODH/ACS) complex (reaction CODH in Fig. 2A), consistent with previous studies [35, 36, 48]. The remaining acetyl-CoA was utilized for carbon metabolism via gluconeogenesis and the incomplete oxidative TCA cycle (reactions POR2 and CS in Fig. 2A).

**Figure 1.**
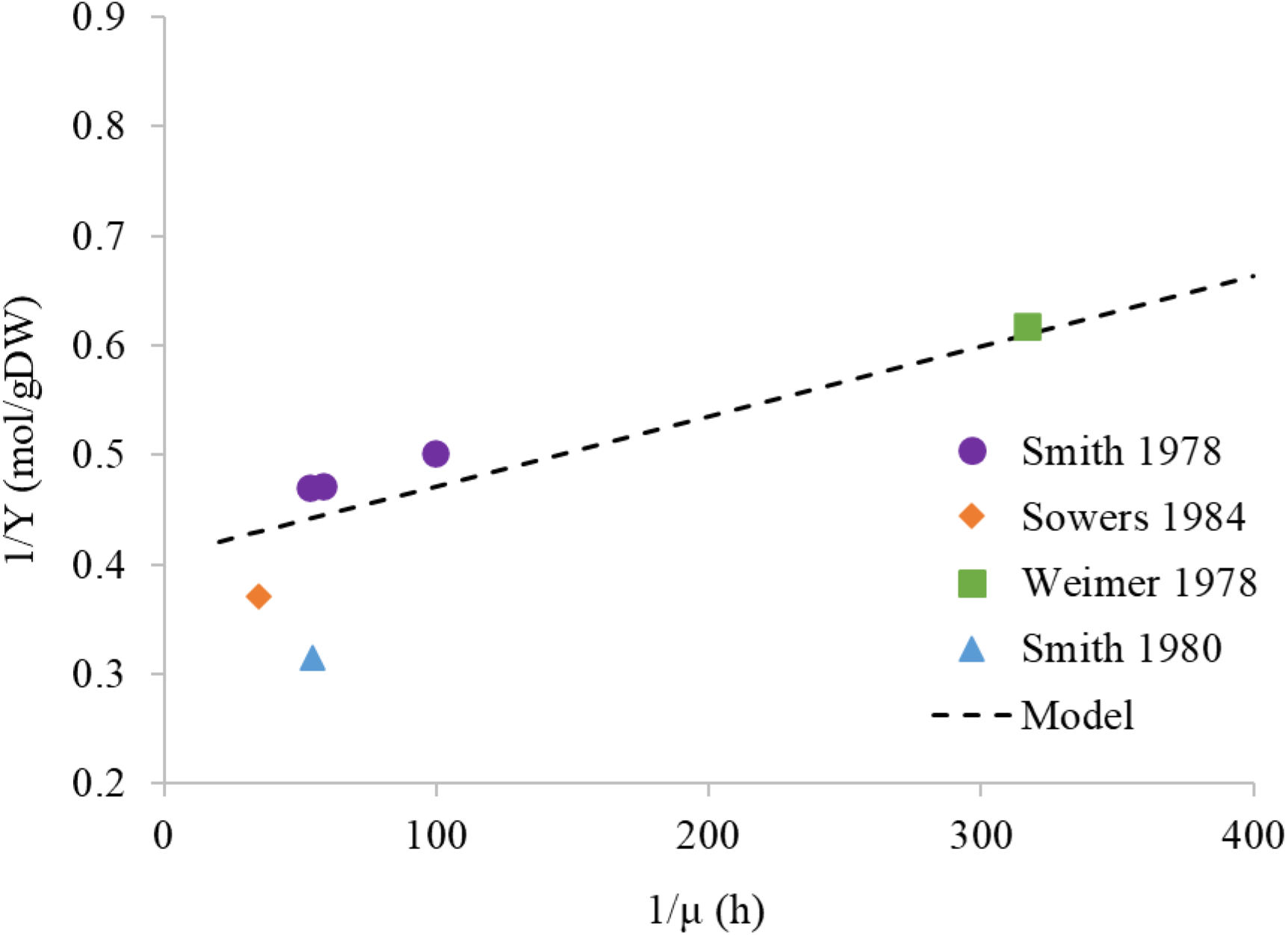
Model validation of *M. barkeri* under aceticlastic methanogenesis. Experimental data were obtained and compiled from various literatures [35, 47–49].

**Figure 2.**
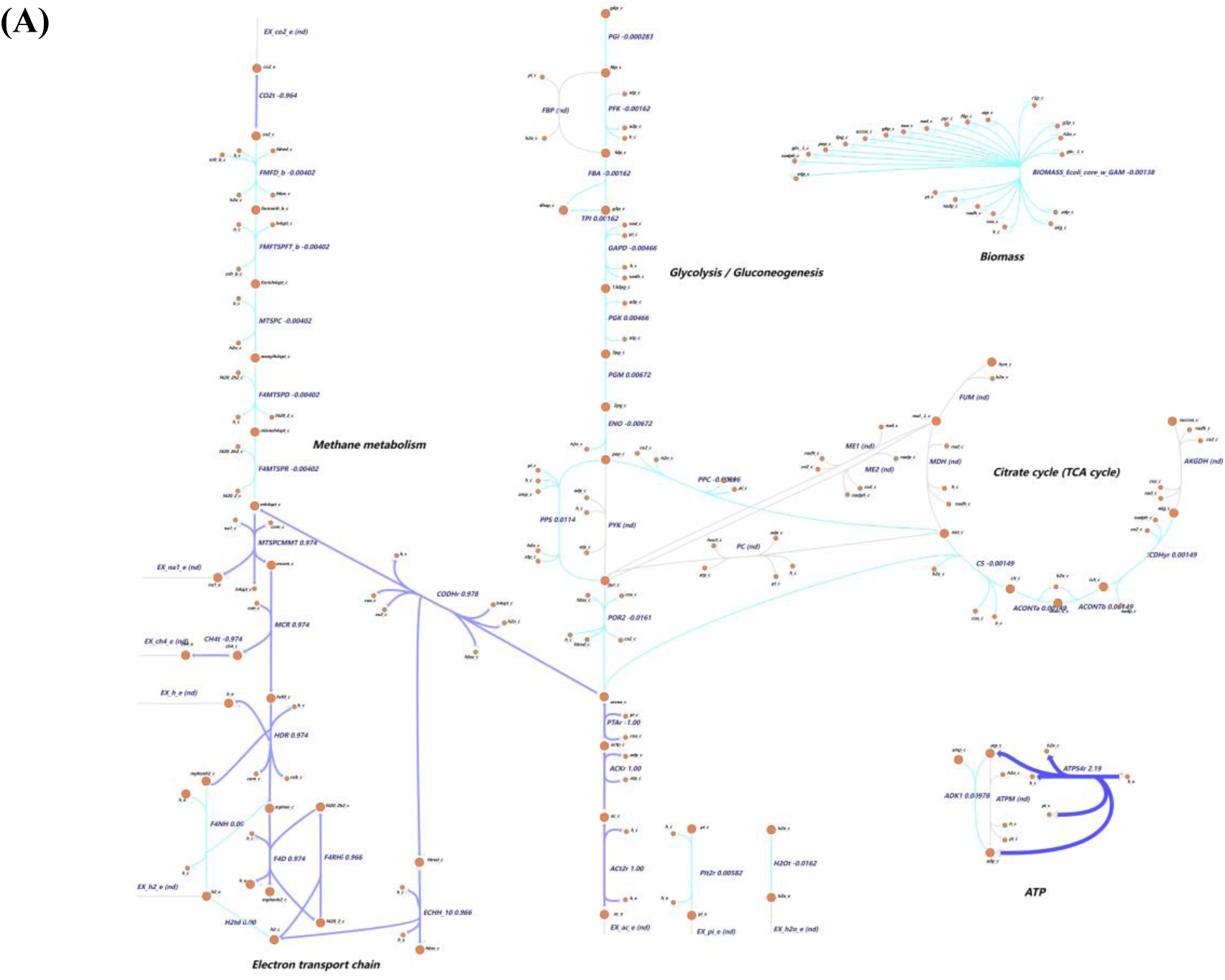
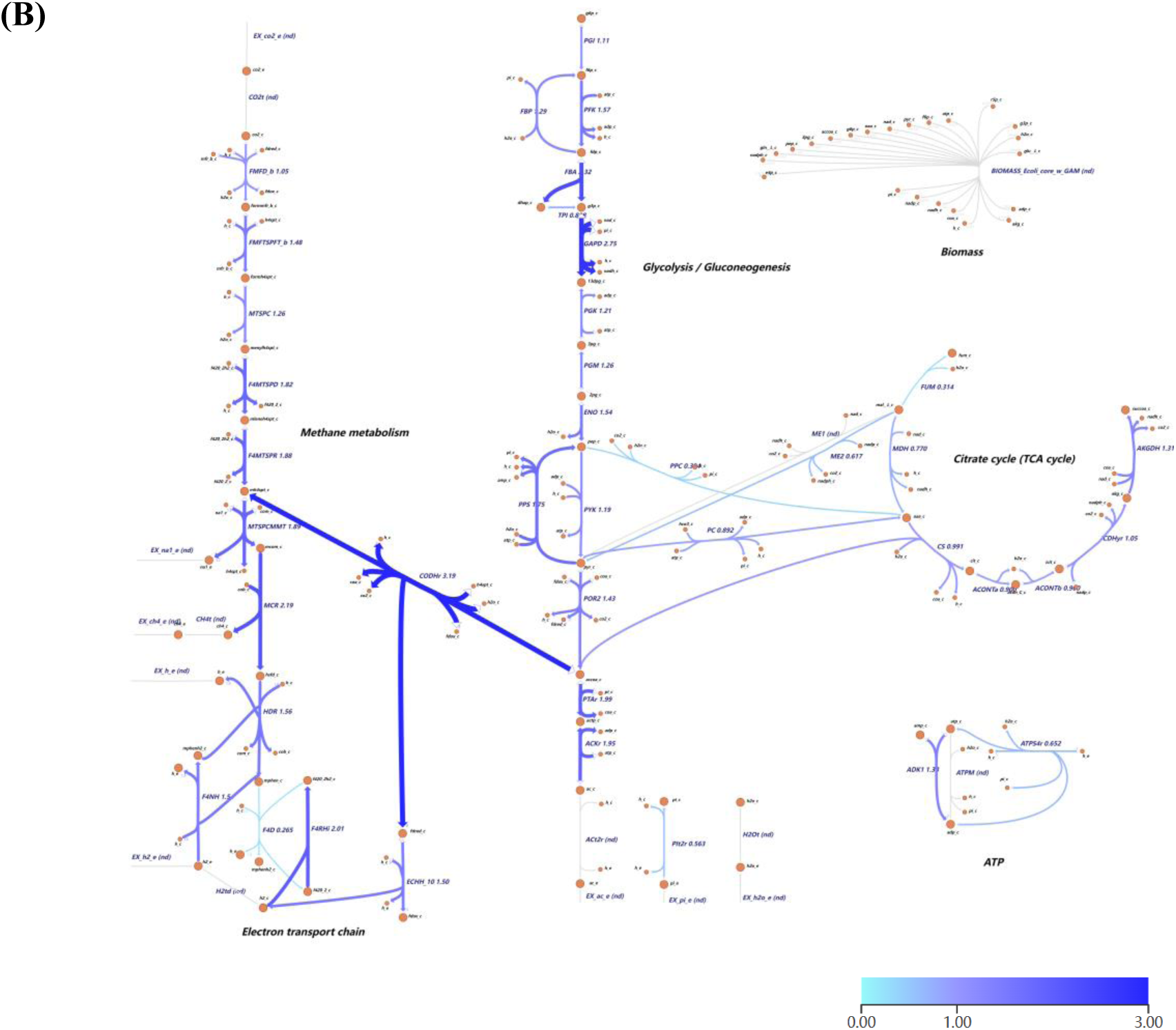
(A) Flux distribution and (B) gene expression levels visualized on central carbon and energy metabolism network of *M. barkeri* under aceticlastic methanogenesis. Orange circles represent metabolites, blue arrows represent active reactions carrying flux, and grey arrows represent unused reactions. Reaction name abbreviations are uppercase and metabolite name abbreviations are lowercase. The acetate uptake rate is set to 1 mmol/gDW/hr (millimoles per gram dry cell weight per hour, the default flux units used in the COBRA Toolbox), the flux value of each reaction is next to the reaction name and indicated by the shades of colour. The gene expression levels are relative to the median value for easy comparison.

Interestingly, the CO_2_ reduction pathway was predicted to actively engage in aceticlastic methanogenesis, which is contrary to previously held assumptions [16, 17]. Previously, it was assumed that acetyl-CoA was split by CODH into a carbonyl group, which would be oxidized into CO_2_ while generating reduced ferredoxin (Fd_red_), and a methyl group, which would be reduced into methane using electrons derived from Fd_red_ via the electron transport chain (Fig. 3A), thus resulting in no net production of Fd_red_ [16, 17]. However, the present study challenged that view by revealing that a minor fraction of acetyl-CoA was utilized for anabolism through the reversible pyruvate: ferredoxin oxidoreductase (POR), requiring an additional source of Fd_red_ for the reduction of acetyl-CoA into pyruvate. This study thus predicted that the CO_2_ reduction pathway operated in a reverse, oxidative direction, generating Fd_red_ to reduce acetyl-CoA and F_420_H_2_ to fuel the electron transport chain (Fig. 3B). This aligns with the findings that a portion of acetate methyl groups are oxidized into CO_2_ during aceticlastic methanogenesis [51]. More importantly, this result is supported by evidence from gene deletion studies [52, 53]. Specifically, disruptions in the CO_2_ reduction pathway—such as deletion of the *mch* gene [52] or the *mtr* operon [53]—significantly impair aceticlastic methanogenesis (Fig. 3A) and inhibit growth on acetate. Transcriptomic analysis of *M. barkeri* cells further supports this finding, as genes related to CO_2_ reduction pathway were highly expressed under aceticlastic methanogenesis (Fig. 2B) [20].

**Figure 3.**
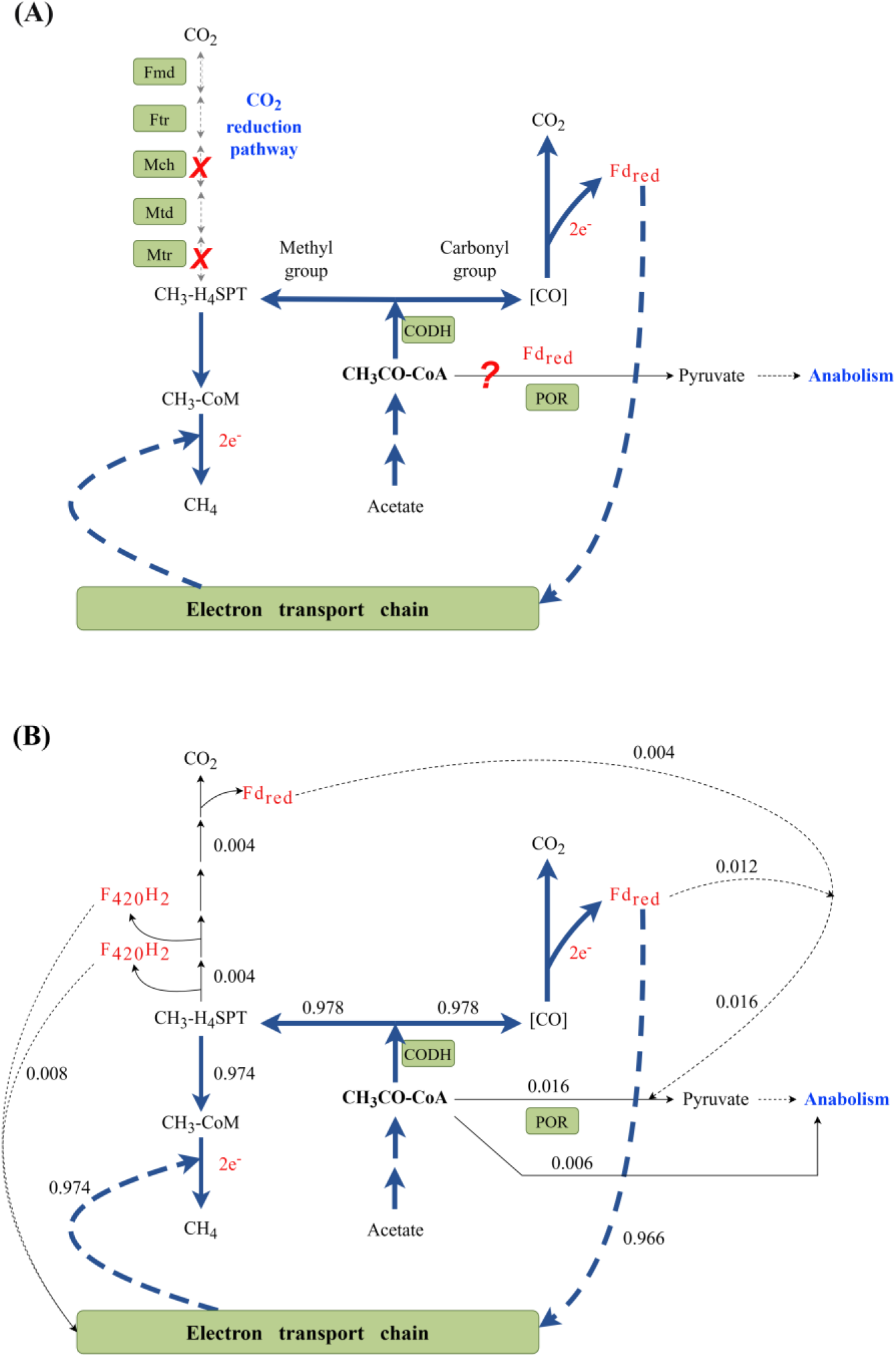
A schematic comparison of the overall electron transfer in aceticlastic methanogenesis pathway: (A) previous assumption and (B) predictions from this study. Gene deletion studies demonstrated that Mch and Mtr in the CO_2_ reduction pathway were essential for growth on acetate, consistent with the model predictions. Abbreviations: CODH, carbon monoxide dehydrogenase; Fmd, formylmethanofuran dehydrogenase; Ftr, formyltransferase; Mch, methenyl-H_4_SPT cyclohydrolase; Mtd, F_420_-dependent methylene-H_4_SPT dehydrogenase; Mer, methylene-H_4_SPT reductase; Mtr, methyl-H_4_SPT: CoM methyltransferase; POR, pyruvate: ferredoxin oxidoreductase.

This study also elucidated the functional role of electron transport chains during aceticlastic methanogenesis. *M. barkeri* conserves energy efficiently through an electron transport chain that coupled acetyl-CoA cleavage with the generation of a transmembrane ion gradient [15]. During aceticlastic methanogenesis, the electron carrier Fd_red_, produced from acetyl-CoA cleavage via CODH, donated electrons to the energy-conserving hydrogenase (Ech). This process (reactions CODH and ECHH in Fig. 2A) coupled Fd_red_ oxidation with H_2_ generation and proton translocation, contributing to ion gradient generation. Based on our constructed model, two distinct pathways, H_2_ cycling pathway and F_420_H_2_-dependent pathway, were identified as capable of transferring electrons from H_2_ to heterodisulfide reductase (Hdr), resulting in the translocation of six protons for ATP synthesis (Fig. 4). The H_2_ cycling pathway included H_2_ generation via Ech (reaction ECHH in Fig. 2A), H_2_ diffusion, H_2_ oxidation and methanophenazine (MP) reduction via viologen-reducing hydrogenase two (Vht, reaction F4RH), and MPH_2_ oxidation coupled with CoM-S-S-CoB heterodisulfide reduction by Hdr (reaction HDR). Alternatively, the F_420_H_2_-dependent pathway utilized the electron carrier coenzyme F_420_H_2_, encompassing H_2_ generation via Ech, H_2_ oxidation and F_420_ reduction via F_420_-reducing hydrogenase (Frh, reaction F4RH), F_420_H_2_ oxidation and MP reduction via Fpo (reaction F4D), and Hdr. Biochemical studies suggest that H_2_ cycling is the preferred pathway for electron transfer during growth on methanol, with the F_420_H_2_-dependent pathway serving as a supplementary role in absence of H_2_ cycling [17, 54, 55]. Nevertheless, the specific contributions of these two pathways during aceticlastic methanogenesis were not well-defined.

**Figure 4.**
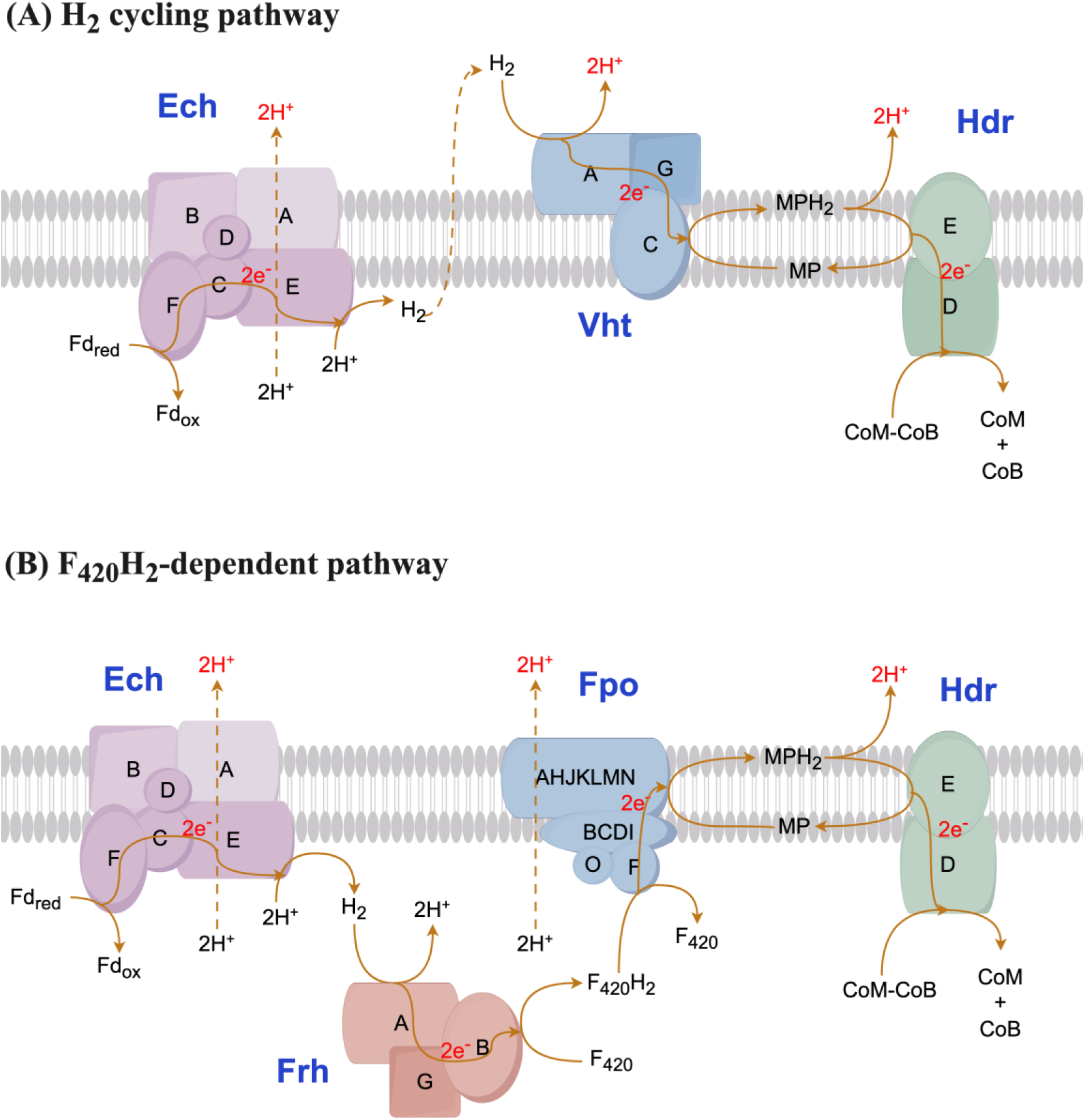
Electron transport chain consisting of two different electron transfer pathways: (A) H_2_ cycling pathway, including energy-conserving hydrogenases (Ech), methanophenazine-reducing Vht hydrogenase, and heterodisulfide reductase (Hdr); and (B) F_420_H_2_-dependent pathway, including Ech, F_420_-reducing hydrogenase (Frh), F_420_: phenazine oxidoreductase (Fpo), and Hdr.

FBA analysis in this study identified the F_420_H_2_-dependent pathway as the primary route for electron transfer during aceticlastic methanogenesis (reactions F4RH and F4D in Fig. 2A). Transcriptomic data supported this finding, showing higher expression levels of genes encoding the F_420_H_2_-dependent pathway (F4RH and F4D in Figure 2B) compared to those of the H_2_ cycling pathway (F4NH in Fig. 2B) [20]. Moreover, the high intracellular coenzyme F_420_ content further confirmed its active role as an electron carrier during aceticlastic methanogenesis [56, 57]. Nevertheless, FVA analysis was conducted to explore the viable range of reaction fluxes, and the results showed that electron transfer through H_2_ cycling pathway could also yield a feasible solution. Experimental evidence demonstrated that H_2_ could serve as a reservoir of reducing equivalents at high acetate concentration, suggesting a potential role for the H_2_ cycling pathway during aceticlastic methanogenesis [58]. In summary, the F_420_H_2_-dependent pathway is the dominant electron transfer pathway during aceticlastic methanogenesis, with the H_2_ cycling pathway potentially serving as a supplementary pathway under certain conditions, such as elevated acetate concentrations.

### 3.2. Metabolic modelling of DIET-based methanogenesis

A previous transcriptomic study explored the gene expression patterns of *M. barkeri* under both DIET and MIET conditions, and a putative MP-reducing enzyme was proposed to be responsible for extracellular electron uptake during DIET-based methanogenesis [18]. However, bioelectrochemical studies indicated that hydrogenases actively engaged in the uptake of extracellular cathodic electrons, resulting in the formation of H_2_ [59, 60]. Despite this, it remained unclear whether H_2_ was abiotically generated at the cathode and diffused into *M. barkeri* as an electron mediator (i.e., in MIET mode), or whether hydrogenases directly captured electrons from the cathode in DIET mode and then converted them into H_2_. Subsequent studies provided evidence supporting the direct uptake of cathodic electrons by hydrogenases, effectively ruling out the use of H_2_ as a diffusible electron mediator in MIET mode [61].

Building on these findings, the current study proposed that the transmembrane Vht hydrogenase served as the interface for direct electrons uptake (Fig. 5). The model predicted that protons released from *G. metallireducens* would combine with electrons in an equimolar ratio to form H_2_ via Vht hydrogenase, ensuring balanced electron and proton flux. This H_2_, along with H_2_ generated from Ech hydrogenase using electrons from acetate cleavage in the form of Fd_red_, was predicted to be oxidized by Frh to produce F_420_H_2_ (Fig. 5). This F_420_H_2_ was then utilized both for MP reduction via Fpo and for the CO_2_ reduction pathway. Collectively, these processes were anticipated to yield a methane-to-ethanol ratio of 1.44, which closely matched the experimentally determined value of 1.5 [8], demonstrating the predictive accuracy of current model. This model-based updated electron transfer mechanism provides a theoretical framework for understanding DIET-based methanogenesis, which is challenging to be experimentally investigated due to biological and technical limitations.

**Figure 5.**
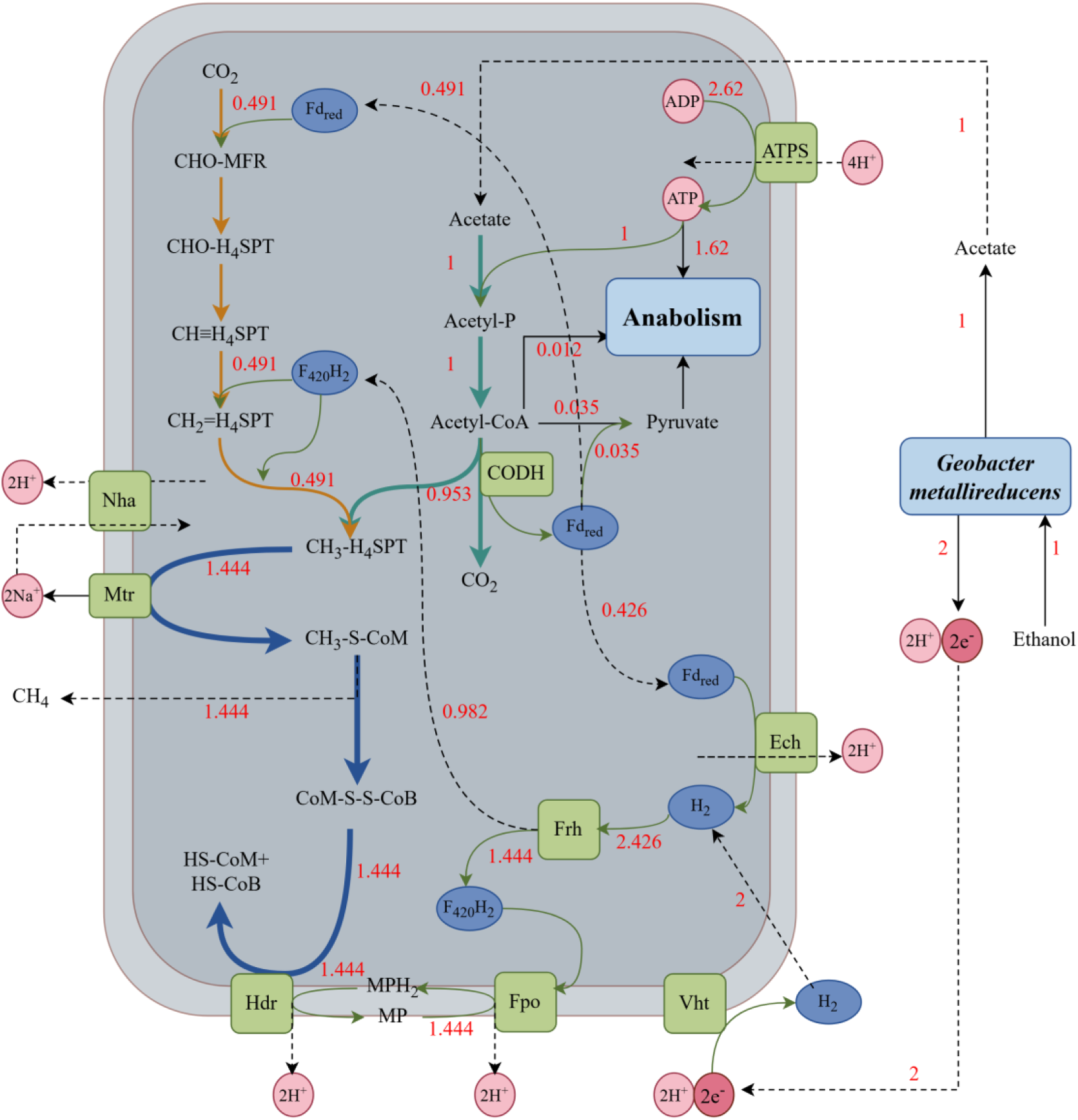
Flux distribution of energy metabolism in *M. barkeri* under DIET-based methanogenesis. The ethanol uptake rate is set to 1 mmol/gDW/hr. Abbreviations: ATPS, ATP synthase; Ech, energy-conserving hydrogenase; F_420_H_2_, reduced coenzyme F_420_; Fd_red_, reduced ferredoxin; Fpo, F_420_: phenazine oxidoreductase; Frh, F_420_-reducing hydrogenase; Hdr, heterodisulfide reductase; MPH_2_, reduced methanophenazine; Mtr, methyl-H_4_SPT: CoM methyltransferase; Nha, Na^+^/H^+^antiporter; Vht, viologen-reducing hydrogenase two.

This study also re-evaluated the previously proposed metabolic pathway for DIET-based methanogenesis [18] using metabolic modelling and compared it with our current model. As depicted in Fig. 6A and 6B, the primary difference lied in the electron transfer chain. The earlier study proposed that Fpo operated in the MPH_2_ oxidation and F_420_ reduction direction (reaction F4D in Fig. 6B, indicated by a negative flux value), in contrast to the flux direction predicted by the current model (Fig. 6A, positive flux value). In the previous model, reduced coenzyme F_420_H_2_ was then proposed to supply electrons for Fd_red_ generation via the electron-bifurcating HdrABC [18]. However, this assumption did not account for the role of aceticlastic methanogenesis, which occurs concurrently with CO_2_ reduction during DIET [8, 18]. Our model predicted that aceticlastic methanogenesis pathway could generate sufficient Fd_red_ (0.953 Fd_red_ molecules per ethanol uptake) via CODH to meet the requirement of CO_2_ reduction (0.491 Fd_red_ molecules per ethanol uptake) (Fig. 5 and Fig. 6B), thereby challenging the previously assumed necessity of HdrABC for Fd_red_ regeneration. In addition, Vht (reaction F4NH) was not predicted to be functional based on the previous assumption (Fig. 6B), as a putative transmembrane enzyme was proposed for extracellular electron uptake and MP reduction to MPH_2_ [18], conflicting with the functional role of Vht. However, genes encoding Vht were actively expressed based on the transcriptomic study (visualized as reaction F4NH in Fig. 6D), and bioelectrochemical studies demonstrated that hydrogenases play an active role in DIET [59, 61]. By considering aceticlastic methanogenesis and the active role of Vht hydrogenase, the model developed in this study provides a more reasonable representation of DIET-based methanogenesis.

**Figure 6.**
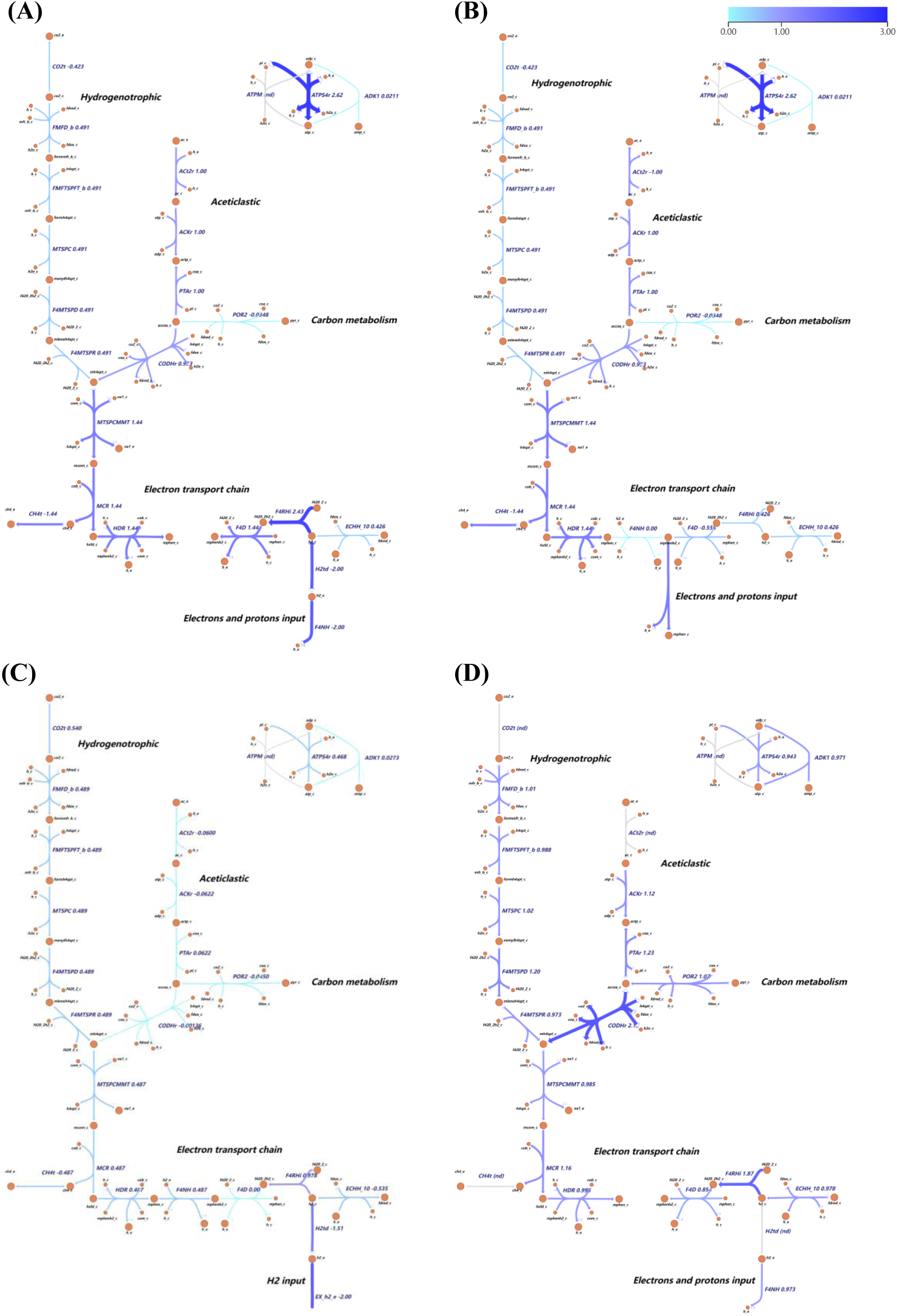
Comparison of flux distribution of energy metabolic pathway: (A) DIET-based methanogenesis predicted in this study; (B) DIET-based methanogenesis based on previous assumption; (C) MIET-based hydrogenotrophic methanogenesis. Gene expression levels under DIET condition was visualized in (D).

Metabolic modelling of MIET-based hydrogenotrophic methanogenesis was also performed and compared with DIET-based methanogenesis (Fig. 6C). Under MIET conditions, only the CO_2_ reduction pathway was active, with acetate serving solely an anaplerotic role in biomass synthesis, as supported by experimental data [8]. As illustrated in Fig. 6A and 6C, the major difference in electron transport between MIET and DIET conditions was related to Fpo (reaction F4D), which was inactive under MIET condition, consistent with previous transcriptomic analysis [18]. Overall, the developed metabolic model in this study effectively captured several transcriptomic features under both DIET and MIET conditions.

### 3.3. Gene expression data and metabolic flux

The weak correlation between gene expression levels and protein abundance presents challenges in deriving biologically meaningful insights from high-throughput transcriptomic data [21, 25, 62]. Metabolic models offer a solution by taking into account network topology, constraints, and interactions that are not captured by gene expression data alone. Consequently, integrating gene expression data with metabolic models has become an area of active research, with several methods being proposed [27]. One widely used approach is the threshold-based method, which applies thresholds to determines the activation status of gene-associated reactions (GAR), by turning the reaction “on” if its expression level is above the threshold [26], or by categorizing the GAR into one of three activity levels: low, medium, or high [27]. The underlying presumption of this approach is that gene expression levels can be utilized to estimate metabolic fluxes with a reasonable degree of accuracy.

However, as shown in Table 1, gene expression data exhibited a weak correlation with model-predicted metabolic flux across all methanogenic conditions, with correlation coefficients below 0.3. This aligns with a previous study reporting significant increases (10-fold) in metabolic flux despite unchanged or reduced gene expression [63]. Violin plots comparison of metabolic model predicted fluxes and gene expression data revealed distinct distributions between these two datasets under different growth conditions (Fig.7). Additionally, gene expression data for DIET condition collected from different studies (DIET and DIET-2) also showed different distributions (Fig.7). Distribution of flux data and expression levels under aceticlastic condition were more elongated, indicating a wider variance. When a threshold-based method was applied to optimize the concordance between gene expression levels and metabolic fluxes (indicated as “Best possible” in Fig. 8) under this condition, 20% of the gene-associated reactions (GAR) displayed active flux while lacking expression, as expression levels below the threshold were falsely turned off. On the other hand, when no threshold was applied (“Entire model” in Fig. 8), all GARs were considered active based on gene expression, but only 42% exhibited active flux, highlighting that gene expression is necessary but not sufficient for enzyme activity. In contrast, under MIET conditions, applying or not applying a threshold made no significant difference (“Entire model” and “Best possible” in Fig. S3). This was supported by the similar and concentrated distribution of these two datasets (Fig. 7). The validity of the threshold-based method depended on the distribution of gene expression data and its correlation with metabolic flux, therefore, this method was not adopted in this study. To avoid excluding essential reactions with low expression levels, a less restrictive strategy was adopted in this study by simply integrating the gene expression data onto metabolic reactions in the model without implementing any thresholds. Consequently, reactions were activated if the corresponding genes were expressed, and deactivated when the corresponding genes were not expressed, ensuring that essential low-expression reactions were not overlooked.

**Figure 7.**
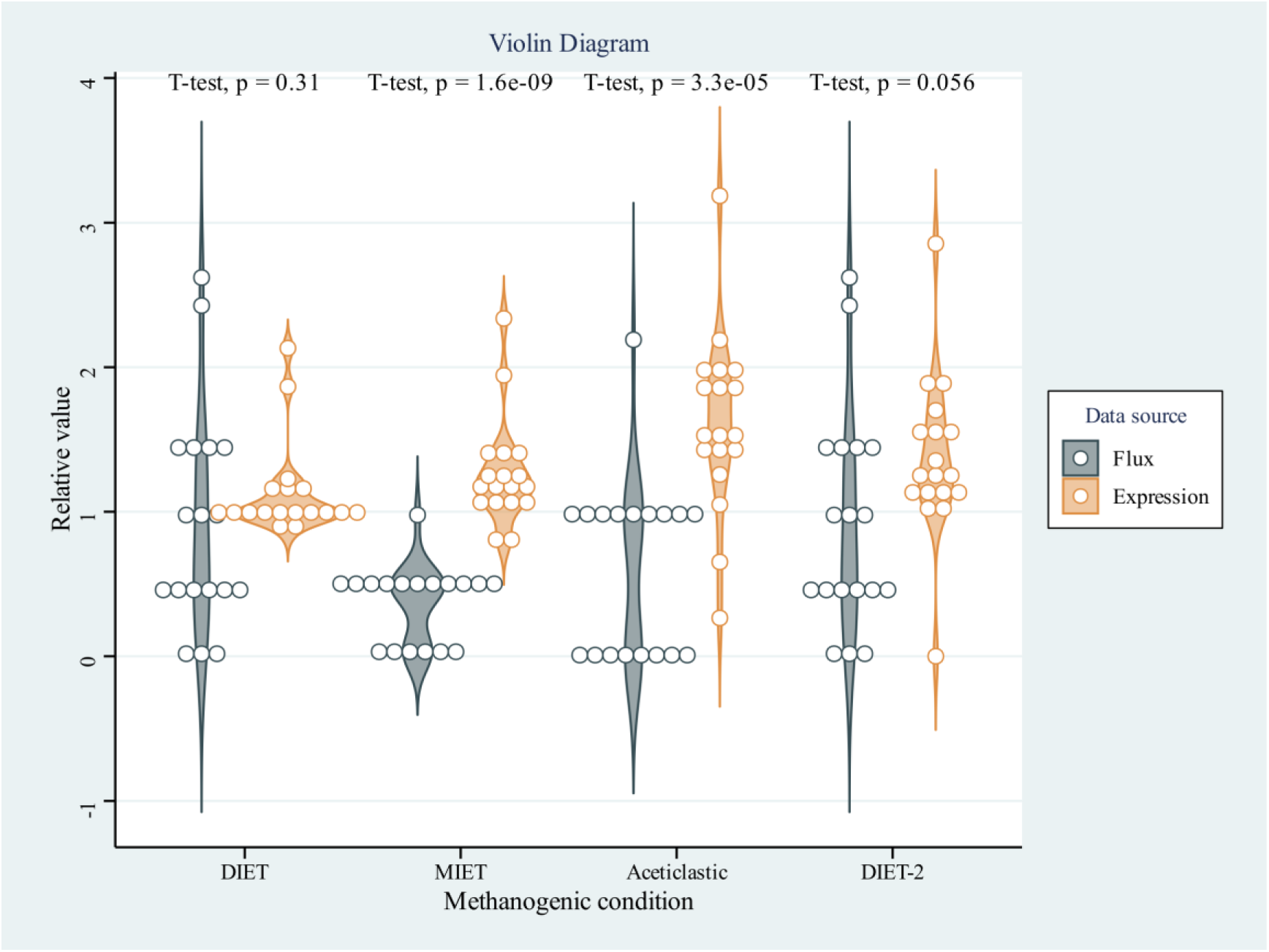
Distribution of metabolic fluxes and gene expression data under different conditions of methanogenesis. Expression data was relative to their median value for easy comparison.

**Figure 8.**
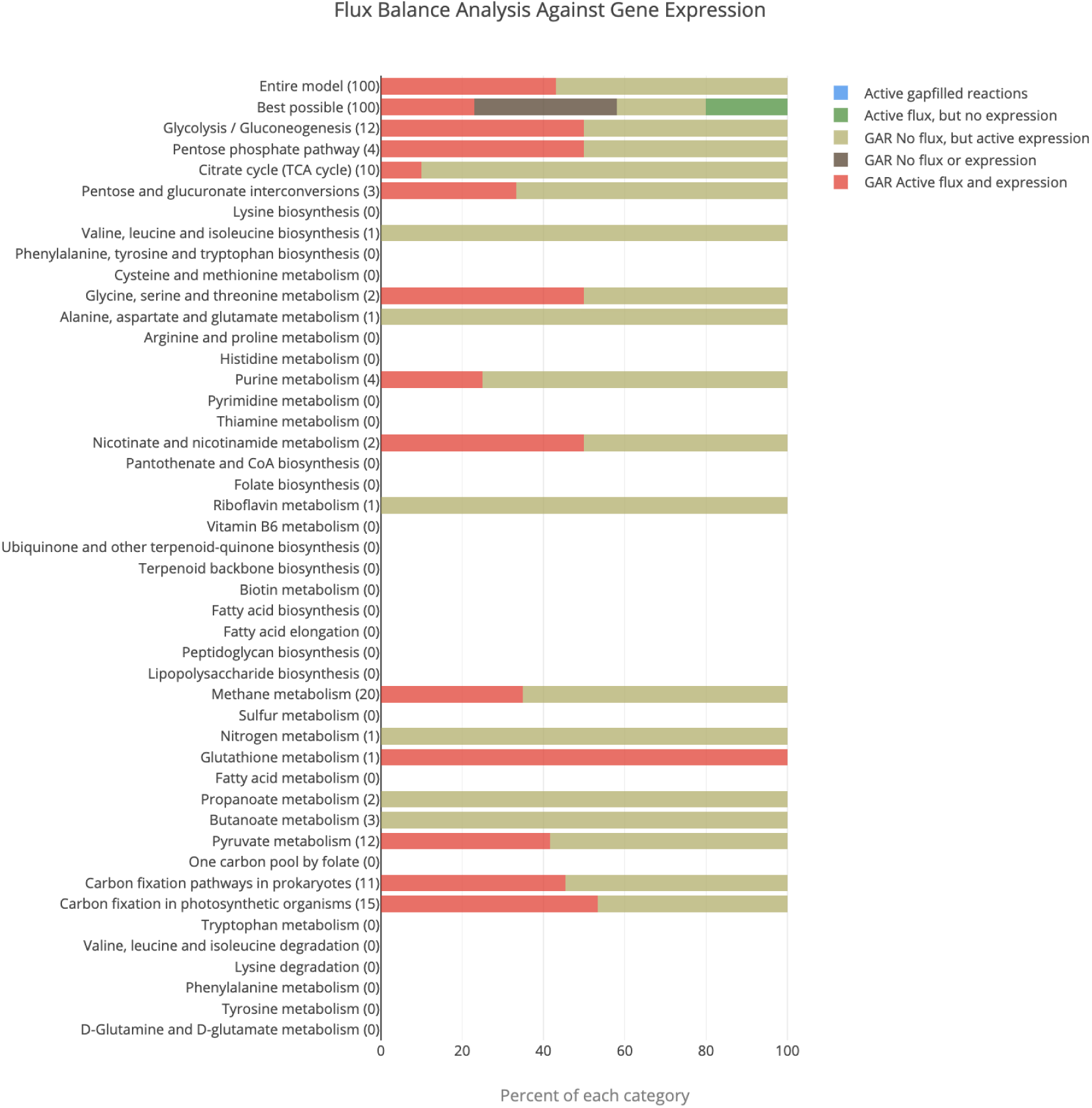
Pathway-based comparison of metabolic flux with gene expression during aceticlastic methanogenesis. “Best possible” means a threshold was applied to maximize the agreement between gene expression and metabolic flux, while “Entire model” donates default setting without threshold.

**Table 1.** Pearson correlation coefficient (r) between gene expression data and metabolic flux.

| Methanogenic condition | r | Accession number | Expression data source |
| --- | --- | --- | --- |
| DIET | 0.282 | SRX4966420, SRX4966418, SRX4966417* | [17] |
| DIET | 0.063 | SRX14923653 | [9] |
| MIET | 0.058 | SRX4966422, SRX4966421, SRX4966419 | [17] |
| Aceticlastic | 0.017 | SRX5544684 | [19] |

\* Publicly available expression data under different growth conditions was attracted from NCBI SRA database with corresponding accession number, and average value was computed for triplicate samples.

This integrative approach was powerful for elucidating the complex mechanisms governing cellular metabolism in *Methanosarcina*. By contextualizing gene expression within the metabolic network, more precise interpretations of the functional implications of gene activity were achieved. For instance, a previous transcriptomic study comparing DIET and MIET conditions proposed an electron transfer pathway involving Fpo and HdrABC based on their high expression levels [18]. However, the metabolic model developed in this study demonstrated that CODH could generate sufficient Fd_red_ for CO_2_ reduction (Fig. 5), thereby challenging the previously assumed necessity of HdrABC for the regeneration of Fd_red_. On the other hand, gene expression data provided the condition-specific insights that are often absent in generic models. For instance, the active express of genes encoding the CO_2_ reduction pathway indicated its functionality under aceticlastic condition. Incorporating this gene expression data into the model activated the functionality of the CO_2_ reduction pathway and uncovered its role in in regenerating Fd_red_. Consequently, gene expression data and metabolic models complement each other, offering a more comprehensive understanding of the electron transfer mechanisms of *M. barkeri* under different methanogenic conditions.

## 4. Conclusions

Linking differential gene expression to functional changes is challenging, primarily due to the weak correlation between gene expression data and protein levels. This study introduced an efficient method for deriving biologically meaningful insights from the growing volume of transcriptomic data by integrating it into the metabolic model of *Methanosarcina barkeri*. Model predictions revealed that the CO_2_ reduction pathway operates in the oxidative direction during aceticlastic methanogenesis, a finding supported by gene deletion studies and in contrast to previous assumptions. Furthermore, the integrated metabolic model effectively captured key transcriptomic features under both DIET- and MIET-based methanogenesis, highlighting the crucial role of the transmembrane hydrogenase Vht in electron uptake under DIET condition. A comparison of gene expression data and model-predicted metabolic flux revealed a weak correlation across all conditions, but they can complement each other to obtain a holistic view of metabolic functions. This approach provides more comprehensive insights into the electron transfer mechanisms of *M. barkeri* under different methanogenic conditions, advancing our understanding of methanogenesis and providing a basis for developing strategies to manipulate this process.

## Acknowledgments

This research was supported by the Excellent Young Scientists Fund of National Natural Science Foundation of China (grant no. 52222008), the Science and Technology Development Fund, Macau SAR (China) (Nos. 0026/2022/A1; 0103/2024/AMJ; 0030/2024/AGJ), Shenzhen-Hong Kong-Macau Science and Technology Project (grant no. EF2023-00072-FST), Research Committee of University of Macau (grant nos. MYRG-GRG2023-00062-FST-UMDF; MYRG2022-00041-FST), the Research Grants Council of Hong Kong Special Administrative Region, China (grant no. T21-604/19-R), and the Hong Kong Innovation and Technology Commission (grant no. ITC-CNERC14EG03).

## Abbreviations

ACK: acetate kinase
CODH/ACS: carbon monoxide dehydrogenase/acetyl-CoA synthase
DIET: direct interspecies electron transfer
Ech: energy-conserving hydrogenase
FBA: flux balance analysis
F_420_H_2_: reduced coenzyme F_420_
Fd_red_: reduced ferredoxin
Fmd: formylmethanofuran dehydrogenase
Fpo: F_420_: phenazine oxidoreductase
Frh: F_420_-reducing hydrogenase
Ftr: formyltransferase
FVA: flux variability analysis
GAR: gene associated reactions
GEMs: genome-scale metabolic models
Hdr: heterodisulfide reductase
Mch: methenyl-H_4_SPT cyclohydrolase
Mer: methylene-H_4_SPT reductase
MIET: mediated interspecies electron transfer
MP: methanophenazine
MPH_2_: reduced methanophenazine
Mtd: F_420_-dependent methylene-H_4_SPT dehydrogenase
Mtr: methyl-H_4_SPT:CoM methyl-transferase
Nha: Na^+^/H^+^ antiporter
POR: pyruvate: ferredoxin oxidoreductase
PTA: phosphotransacetylase
TCA cycle: tricarboxylic acid cycle
Vht: viologen-reducing hydrogenase two

